# The new *SH3_T* domain increases the structural and functional variability among SH3b-like CBDs from staphylococcal phage endolysins

**DOI:** 10.1101/2024.04.22.590666

**Authors:** Roberto Vázquez, Diana Gutiérrez, Dennis Grimon, Lucía Fernández, Pilar García, Ana Rodríguez, Yves Briers

## Abstract

Endolysins, proteins encoded by phages to lyse their hosts and release their progeny, have evolved to adapt to the structural features of each host. The endolysins from *Staphylococcus*-infecting phages typically feature complex architectures with two enzymatically active domains (EADs) and one cell wall-binding domain (CBD) belonging to the bacterial SH3 (SH3b) superfamily. This study focuses on three SH3b-like CBDs from exemplary staphylococcal phage endolysins (LysRODI, LysC1C, and LysIPLA5) that were structurally and functionally characterized. While RODI_CBD and C1C_CBD were assigned to the well-known *SH3_5* family, a new family, *SH3b_T*, was identified using the CBD from LysIPLA5 as a model. GFP-fused CBDs were created to assess their differential binding to a collection of staphylococcal strains. IPLA5_CBD showed enhanced binding to *Staphylococcus epidermidis*, while RODI_CBD and C1C_CBD exhibited distinct binding profiles, with RODI_CBD targeting *Staphylococcus aureus* specifically and C1C_CBD displaying broad binding. Sequence comparisons suggested that a few differences in key amino acids could be responsible for the latter binding difference. The CBDs modulated the activity spectrum of synthetic EAD-CBD combinations in accordance with the previous binding profiles, but in a manner that was also dependent on the EAD present in the fusion protein. These results serve as a context for the diversity and versatility of SH3b domains in staphylococcal endolysins, providing insights on how (i) the CBDs from this superfamily have diverged to adapt to diverse bacterial ligands in spite of sharing a common fold; and (ii) the evolution of specificity relies on the EAD-CBD combination rather than solely the CBD.

**IMPORTANCE:** Clinical management of bacterial infections is nowadays compromised by the rise in antimicrobial resistance. The development of new antimicrobial therapies with diverse modes of action is therefore of pivotal importance to complement the current standard of care. Phage endolysins are a new class of antibacterial agents based on rapid peptidoglycan degradation. The natural reservoir of phage endolysins offers a practically infinite diversity. This works reveals a broadly spread but still unknown phage endolysin domain targeting staphylococci while providing structural-functional insights that are paramount to understand the evolution of endolysins and how they can be applied as an antimicrobial.

## INTRODUCTION

Endolysins are one of the gene products that dsDNA (bacterio)phages use to release the viral progeny from their host bacterial cells. They contain a catalytic, peptidoglycan-degrading activity, and thus when released to the periplasm via different tightly regulated mechanisms, they provoke bacterial cell lysis by osmotic shock due to the disruption of the peptidoglycan (1). Besides their natural key role in the phage infection cycle, endolysins have sparked interest since they can be purposed as alternative antimicrobial agents (2–4). Due to the escalating burden of antibiotic resistance among clinically relevant bacteria (5), the discovery and development of novel antimicrobials is one of the current main scientific priorities set up by many healthcare authorities and international organizations (6, 7). Recombinantly produced endolysins have been extensively shown to be effective as exogenous antibacterial agents *in vitro* and *in vivo*, and different clinical trials have been conducted (8–10). Besides their lesser probability to cause resistance in bacteria, the most interesting features of lysins both from an applied and a fundamental perspective are probably their great natural variability and modularity (Criel *et al*, 2021; Vázquez *et al*, 2021). Both characteristics are intertwined, as phages rapidly and dynamically evolve in a modular manner, exchanging functionally autonomous modules of genetic information between each other and with their bacterial hosts (11). At the endolysin level, their modularity means that they may comprise different functional domains: one or more enzymatically active domains (EADs) and a cell wall-binding domain (CBD). Usually, phages that infect Gram-negative hosts bear lysins with a single EAD, whereas those from Gram-positive hosts typically have several domains, at least one of each kind (12, 13). The preferential presence of CBDs in endolysins from a Gram-positive background is hypothetically explained either by (i) the need for a tropism of the enzyme towards its insoluble, non-diffusible substrate (as is the case for many enzymes acting on polymeric substrates (14); (ii) for the endolysin to remain tightly bound to the cell debris of the lysed host thus preventing the killing of neighboring cells that are potential new hosts for the phage progeny; or by a combination of both reasons (15). Importantly, due to their typically high affinity, CBDs are thought to be the main determinant for the observed endolysin specificity, as proven, for example, by domain swapping experiments in endolysins derived from phages infecting *Streptococcus* or *Listeria* (Diez-Martinez *et al*, 2015; Schmelcher *et al*, 2011; Vázquez *et al*, 2017).

The case of endolysins from staphylococcal phages has been extensively studied due to the prominent role of many staphylococcal species in human or animal microbiota and disease (16). For example, *Staphylococcus aureus* is one of the most burdensome human bacterial pathogens globally (17), and *Staphylococcus epidermidis* plus some other so-called coagulase-negative staphylococci are widespread components of the human skin microbiota that are also responsible for nosocomial infections (18). Endolysins from staphylococcal phages have a typical bicatalytic structure (Vázquez *et al.*, 2021), with evolutionarily conserved CBDs belonging to the bacterial SH3 (SH3b) superfamily (19). SH3b (bacterial Src Homology 3) is a superfamily of widespread ligand-binding domains that appear in many bacterial and phage proteins and are also related to homologous ligand-binding domains in other kingdoms (20). The SH3 fold is one of the simplest and oldest ones (21) and its main structural feature is a β-barrel layout usually devoted to a ligand-binding function. The SH3b domains in particular are mainly known to bind cell wall motifs, thus playing a prominent role in cell wall-remodeling enzymes, autolysins and phage endolysins. In the particular case of staphylococcal lysins, the studied CBDs from SH3b have been classified into the *SH3_5* (PF08460) family, and are assumed to specifically bind the peptide moiety of staphylococcal peptidoglycan, including the peptide stem and the peptide cross-link, as recently shown for the *SH3_5* CBD of lysostaphin (22, 23). However, SH3b domains comprise representatives that, while sharing the characteristic SH3-like β-barrel topology, have evolved to recognize a variety of cell wall ligands. For example, the *SH3_5* from *Lactiplantibacillus plantarum* major autolysin Acm2 is a broad-range CBD that recognizes many different peptidoglycan chemotypes (24), the SH3b-like, *PSA_CBD* (PF18341) domain from *Listeria* phage endolysins recognizes serovar-specific motifs at the cell wall teichoic acids (25), and the SH3b CBD from the endolysin of *Bacillus* phage PBC5 binds to the glycan chain (26).

In this work, we aimed at providing insights on the specificity range of SH3b-like CBDs from staphylococcal endolysins and how they impact the antibacterial spectrum of the lysins in which they are inserted. To this end, we focused on three *Staphylococcus* phage endolysins: LysRODI, LysC1C and LysIPLA5 (27, 28). The binding profiles of the selected CBDs were experimentally characterized both as a standalone and in connection with their ability to modulate the activity range of EADs derived from LysRODI and LysC1C. In this way, we expect this work contributes to understand how the structural diversity of staphylococcal CBDs connects to their peptidoglycan-binding function, and how this ability cooperates with intrinsic features of EADs to produce the experimentally observed activity spectra in lysins purposed for exogenous lysis.

## MATERIALS AND METHODS

### Bacterial strains and culture conditions

The staphylococcal strains used in this work (**Table 1**) were grown in tryptic soy broth (TSB) at 37 °C with shaking (200 rpm) or on TSB plates containing 2% (w/v) bacteriological agar. *Escherichia coli* TOP10 was used for cloning and *E. coli* BL21(DE3) for protein expression. *Acinetobacter baumannii* RUH 134 (29) was used as a control strain. All the former Gram-negative bacteria were grown in LB medium at 37 °C with shaking (200 rpm). For the positive selection of pVTEIII or pVTD3 *E. coli* transformants, 100 μg/ml ampicillin or 50 μg/ml kanamycin were used, respectively, together with 5% (w/v) sucrose to negatively select against plasmids lacking insertion (as explained in (30)). 100 μg/ml ampicillin was used to select transformants of vectors based on pET21(a). Bacterial stocks were made by adding 20% v/v glycerol to grown bacterial cultures and were kept at −80°C.

**Table 1.** Staphylococcal strains used in this work.

| Species | Strain | Source |
| --- | --- | --- |
| *S. epidermidis* | F12 | (Delgado et al. 2009) |
|  | B |  |
|  | DG2n |  |
|  | YLIC13 |  |
|  | LO5RB1 |  |
|  | DH3LIk |  |
|  | Z2LDC14 |  |
| *S. aureus* | Sa9 | (García et al. 2009) |
|  | IPLA1 | (Gutiérrez et al. 2012) |
|  | IPLA16 |  |
|  | 15981 | (Valle et al. 2003) |
|  | V329 | (Cucarella et al. 2001) |
|  | MRSAE10 | Pig skin isolate (unpublished) |
| *S. hominis* | ZL31-13 | (Martín et al. 2012) |
| *S. xylosus* | ZL61-2 |  |
| *S. haemolyticus* | ZL89-3 |  |
| *S. gallinarum* | ZL90-5 |  |
| *S. kloosi* | ZL74-2 |  |

### Plasmid construction and DNA manipulation

The sequences encoding LysRODI, LysC1C and LysIPLA5 were codon optimized (GenSmart Codon optimization), synthetized and cloned into a pET21(a) vector (between *NdeI* and *XhoI* restriction sites) by GenScript (Rijswijk, Netherlands). For all other proteins used in this work, the expression vectors were constructed through the VersaTile workflow as described in (30). In brief, each individual domain (EAD, CBD or eGFP) was PCR-amplified from its source plasmid with specific primers including BpiI and BsaI recognition sites at both the 5’ and 3’ end, according to the VersaTile method. A restriction/ligation reaction with BpiI was carried out with these amplicons to insert them into the entry vector pVTEIII (Amp^R^, Suc^S^). The ligation products were subsequently use for transformation of *E. coli* TOP 10 by electroporation and transformants bearing pVTEIII plasmids with the inserted tile (Amp^R^, Suc^R^) were selected on LB plates with ampicillin and sucrose. The TOP10 cells were used as a source for tiles, which were all confirmed by Sanger sequencing (LGC Genomics) and stored at the VersaTile repository of Ghent University. Tile ligation into the destination vector pVTD3 (Kan^R^, Suc^S^) was conducted by setting up restriction/ligation reactions with BsaI and the appropriate tiles from the repository (*e.g.*, eGFP plus RODI_CBD and a 6×His tag). All chimeric coding sequences were designed with a C-terminal 6×His tag for purification unless otherwise stated. Final constructs were used for transformation of *E. coli* BL21(DE3), selecting the transformants with kanamycin and sucrose, and their sequence was verified by Sanger sequencing. A list of the tiles used in this work, including their source NCBI entry and the delineation coordinates can be found in **Table 1**.

**Table 1.**
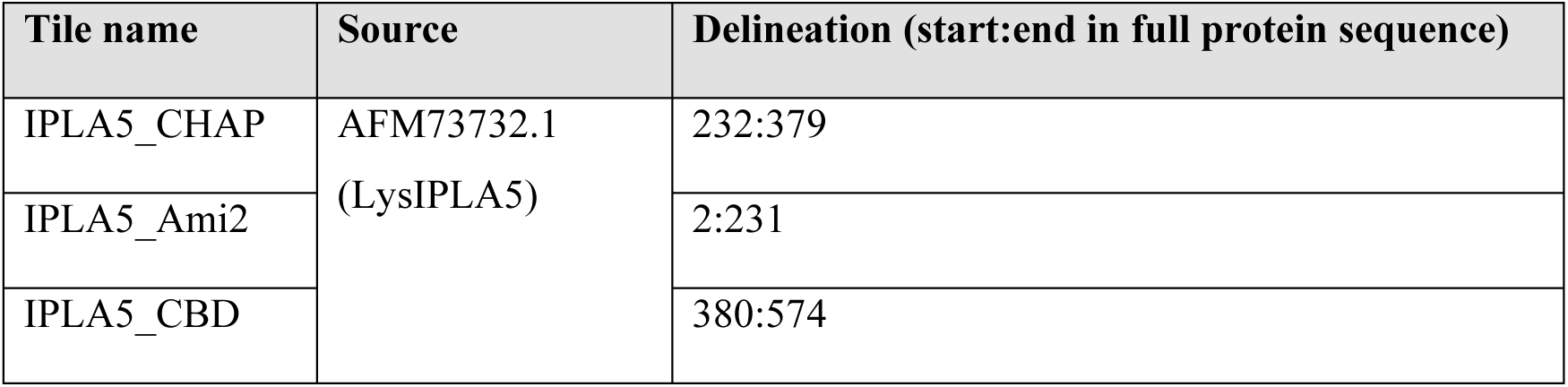
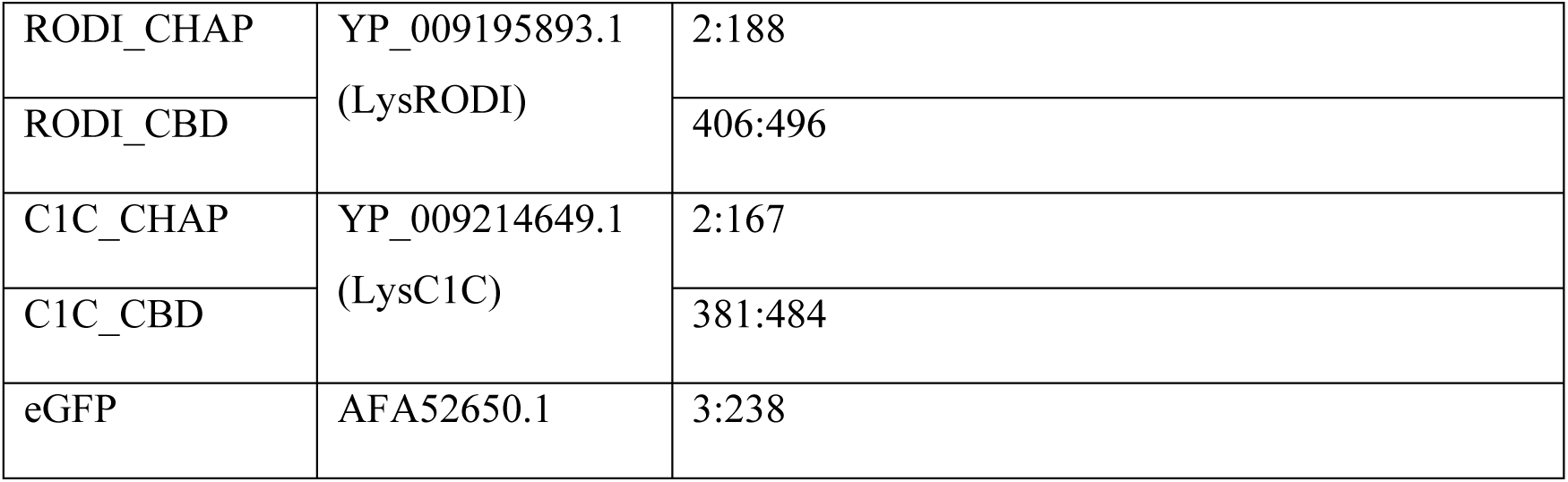
Tiles used in this work.

| Tile name | Source | Delineation (start:end in full protein sequence) |
| --- | --- | --- |
| IPLA5_CHAP | AFM73732.1<br>(LysIPLA5) | 232:379 |
| IPLA5_Ami2 |  | 2:231 |
| IPLA5_CBD |  | 380:574 |
| RODI_CHAP | YP_009195893.1<br>(LysRODI) | 2:188 |
| RODI_CBD |  | 406:496 |
| C1C_CHAP | YP_009214649.1<br>(LysC1C) | 2:167 |
| C1C_CBD |  | 381:484 |
| eGFP | AFA52650.1 | 3:238 |

### Protein expression and purification

The fusion proteins used throughout this work were expressed in *E. coli* BL21(DE3) strains bearing the corresponding pVTD3 or pET21(a) vectors prepared as described in the previous section. Protein expression and purification was performed as previously described (31). After purification, the buffer was exchanged to 50 mM sodium phosphate buffer pH 7.4 using Zeba™ Spin Desalting Columns, 7K MWCO, 5 ml (Thermo Fisher Scientific) following the supplier’s recommendations. Finally, proteins were sterilized by filtration (0.45 μm PES membrane filters, VWR).

Protein concentration was quantified using the Quick Start Bradford Protein assay (BioRad). Relevant information on the proteins used in this work is in **Table 2**.

**Table 2.**
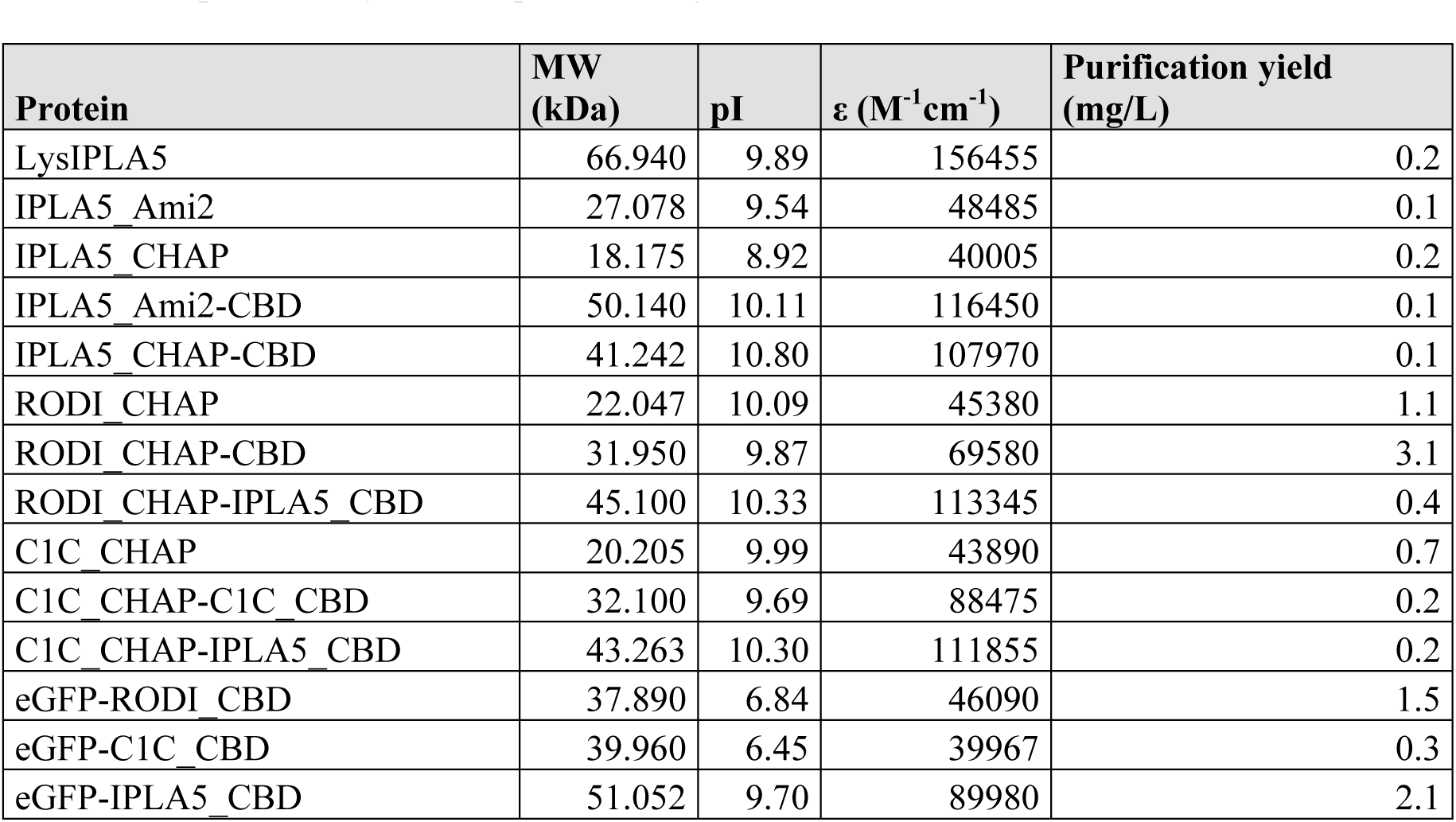
Predicted features (molecular weight, isoelectric point, extinction coefficient) of the proteins used in this work using Expasy ProtParam (https://web.expasy.org/protparam/) along with their experimentally verified purification yield.

### Quantification of bacterial binding

Binding of CBDs to bacterial substrates was measured by recording the fluorescence of eGFP fusions of the different domains. Such fusions were prepared using VersaTile and comprised eGFP at the N-terminal, the CBD at the central and a 6×His tag at C-terminal position. To perform the binding assay, exponential (OD_600_ ≈ 0.5 – 0.6) or stationary phase (OD_600_ ≈ 1– 2) cultures of the strains to be tested were centrifuged (10, 000 × g, 1 min) and the pellets were washed with PBS (137 mM NaCl, 2.7 mM KCl, 10 mM Na_2_HPO_4_, 1.8 mM KH_2_PO_4_, pH 7.4). The bacterial suspensions were adjusted to OD_600_ ≈ 1.0 and dispensed on dark flat bottom 96-well plates (180 µl per well). 20 µl of 20 µM solutions of the eGFP-CBD fusion proteins (or just buffer for bacterial autofluorescence controls) were added to each well and then the plates were incubated for 10 min in the dark at room temperature. Then the plates were centrifuged (1000 × g, 5 min), the supernatants were removed, and the pellets washed once with PBS and finally suspended in 200 µl of PBS. 200 µl of 10 µM fluorescein were added to the plate as internal control to automatically optimize gain, as well as positive fluorescence controls for each eGFP-CBD fusion protein (180 µl PBS plus 20 µl of the 20 µM protein stock solution). Fluorescence was then measured in a TECAN Infinite 200 PRO plate reader (TECAN, Männedorf, Switzerland) with excitation/emission wavelengths of 485 nm and 530 nm, respectively. Fluorescence measurements were then corrected for comparability between proteins by applying a correction factor F_max_/F_prot_ in which F_max_ is the maximum fluorescence recorded and F_prot_ is the fluorescence of each eGFP-CBD at 2 µM. Fluorescence measurements were acquired for three biological replicates.

### Minimum Inhibitory Concentration

The minimum inhibitory concentrations (MICs) of the antimicrobial proteins in this work were determined by the broth microdilution assay as described before (31). The MIC values reported correspond to the mode of three independent biological replicates.

### Bioinformatic Analyses

Two complementary approaches were taken to build a protein sequence dataset of SH3b-like domains related to staphylococcal lysins (**Supplementary Figure S1**). To find representative sequences of the IPLA5_CBD family, termed *SH3b_T*, a phmmer search (32) was conducted against Reference Proteomes restricted to viral taxa (taxid: 10239) and using the first IPLA5_CBD as query (UniProt I6T7G5, from coordinate 380 to 458). The 38 significant hits (*i.e.*, sequences comprising only the *SH3b_T*-like domains) were clustered with CD-HIT and an exclusion cutoff of 97% identity was applied to decrease redundancy (33). A second phmmer iteration was conducted, now against the full UniProt database, using two queries: the first repeat of the LysIPLA5 CBD and the *SH3b_T* domain from A0A499SIE6 (positions 169 to 250), displaying the lowest % identity, in amino acid sequence, with the former (39%). This yielded 366 significant hits. A length cutoff was applied to the sequences (source full proteins > 100 amino acids; query coverage > 60 amino acids) and a second CD-HIT redundancy reduction with 97% identity cutoff was applied, reducing the final set to 59 representatives for which additional metadata (domain predictions for the full-length sequences, bacterial hosts) were mined from UniProt or predicted using hmmscan against the Pfam database. Alternatively, representative domain sequences from the *SH3_5* family were retrieved from PhaLP database of phage lysins (34), restricting the search to endolysins from host genus *Staphylococcus* and domain name *SH3_5* (236 entries total). The CBD sequences were extracted from these entries using the delineating coordinates from the Pfam *SH3_5* predictions stored in PhaLP, and then the same length and % identity cutoffs were imposed to obtain 53 final representative *SH3_5* domain sequences. The same PhaLP-based pipeline was used to retrieve *PSA_CBD* examples (type: “endolysin”, domain name: “*PSA_CBD*”), while the phmmer pipeline was used to retrieve *PBC5_CBD* examples. The first SH3b-like repeat of LysPBC5 (A0A218KCJ1, residues 230 to 279) was used as query for the phmmer searches. The dataset built this way can be accessed as **Supplementary Dataset S1**.

Multiple sequence alignments (MSAs) and phylogenetic analyses were performed and represented in R. The packages ‘msa’ and ‘ggmsa’ were used respectively to build the MSAs (with algorithm ClustalW) and to represent them (35, 36). The phylogenetic trees were built based on the MSAs for which a sequence similarity-based pairwise distance matrix was computed, which was then used as input for building an UPGMA phylogenetic tree and representing it (37–39).

Protein three-dimensional structure predictions were performed using AlphaFold2 algorithm as implemented in ColabFold (40) with default parameters plus Amber relaxation, and models were analyzed and rendered with DeepView (41), which was also used for structural alignments.

### Statistical analyses and experimental data representation

Representation of experimental data was performed using R package ‘ggplot2’ (42). Error bars in bar plots represent the standard deviation of three independent replicates. Additional statistical analyses (Kruskal-Wallis test) were performed in R.

## RESULTS

### LysIPLA5 has an atypical bicatalytic architecture with an SH3b-like C-terminal end

The endolysins from phages phiIPLA-RODI and phiIPLA-C1C (*i.e.*, LysRODI and LysC1C) are examples of the canonical *CHAP* (PF05257):*Amidase_2* (PF01510):*SH3_5* architecture described for a majority of staphylococcal endolysins. The *S. epidermidis* phage vB_SepiS-phiIPLA5, however, bears an endolysin (LysIPLA5) with two predicted EADs, *Amidase_2* and *CHAP*, in the reverse order. In addition, LysIPLA5 has a C-terminus where usually a CBD would be present, but no CBD could be predicted (**Figure 1a**). In a preliminary functional study of LysIPLA5 domains, neither the full protein nor any of its individual domains showed *in vitro* antimicrobial activity against *S. epidermidis* F12, probably due to very low expression yields and subsequently low working concentrations, at least for the full protein (**Supplementary Figure S2**). The C-terminal domain, henceforth referred to as IPLA5_CBD, was nevertheless efficiently expressed and a concentration as high as 34.16 µM was also unable to exert any growth inhibition. On the other hand, whereas ∼6 µM IPLA5_CHAP was also inactive, a MIC (0.75 µM) was achieved when IPLA5_CHAP was fused to the native C-terminal domain IPLA5_CBD. These results initially supported the possibility of IPLA5_CBD being a CBD.

**Figure 1.**
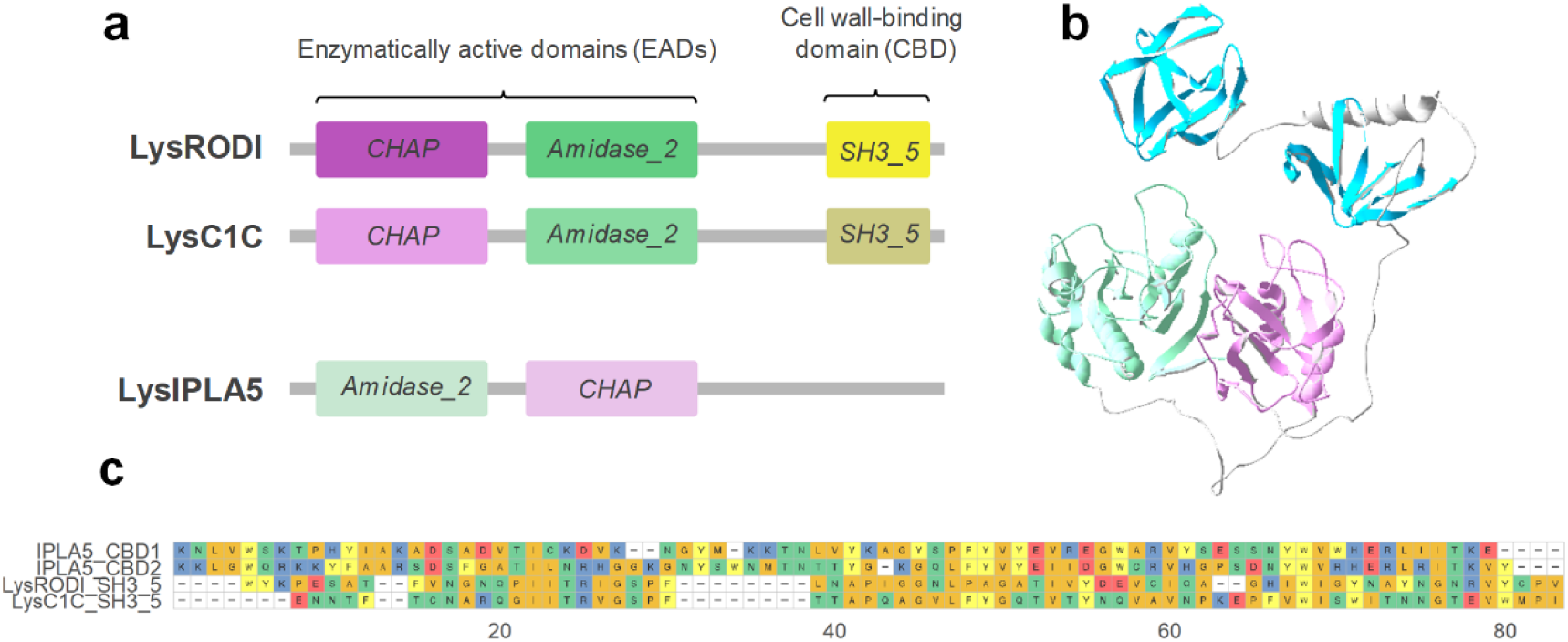
Structural comparison of three staphylococcal endolysins. (a) Schematic representation of the architectures of LysRODI, LysC1C and LysIPLA5 (not in scale). (b) Prediction of LysIPLA5 3D structure (pLDDT = 79.78). (c) Sequence alignment of the repeats making up the CBDs of LysRODI, LysC1C and LysIPLA5.

According to a three-dimensional structure prediction of LysIPLA5, this C-terminal stretch contains two repeats with a fold akin to the typical SH3 β-barrels (**Figure 1b**). The sequences of these repeats substantially differ from the *SH3_5* CBDs of LysRODI and LysC1C (percent identities between 16-21%), while the repeats are 44% identical to each other (**Figure 1c**). Thus, it was concluded that LysIPLA5 bears a C-terminal CBD that belongs to the SH3b superfamily but that constitutes a different family than the usual *SH3_5*, which was then termed *SH3b_T*.

### *SH3b_T* family shows differences in their bacterial distribution with respect to SH3_5

The tree shown in **Figure 2** indicates that the 59 *SH3b_T*-like sequences retrieved by an iterative phmmer search substantially differ in sequence from the 53 established *SH3_5* examples obtained from PhaLP, as they are allocated in different clades and are clearly distinguished in the similarity-based distance matrix heatmap. They also display a marked difference in the bacterial species to which they are associated. In the case of *SH3_5*, they mostly appear in phages whose host is annotated as *S. aureus*, whereas the preferred host for IPLA5_CBD is *S. epidermidis* or other coagulase-negative staphylococci, with only a few exceptions to this rule. In addition, a trend can also be established with respect to the preferred full endolysin architecture: while the preferential architecture for endolysins with an *SH3_5* CBD is the canonical *CHAP*:*Amidase_2*:CBD, the order of the EADs is mostly reversed for lysins with a *IPLA5_CBD*, as in LysIPLA5 itself.

**Figure 2.**
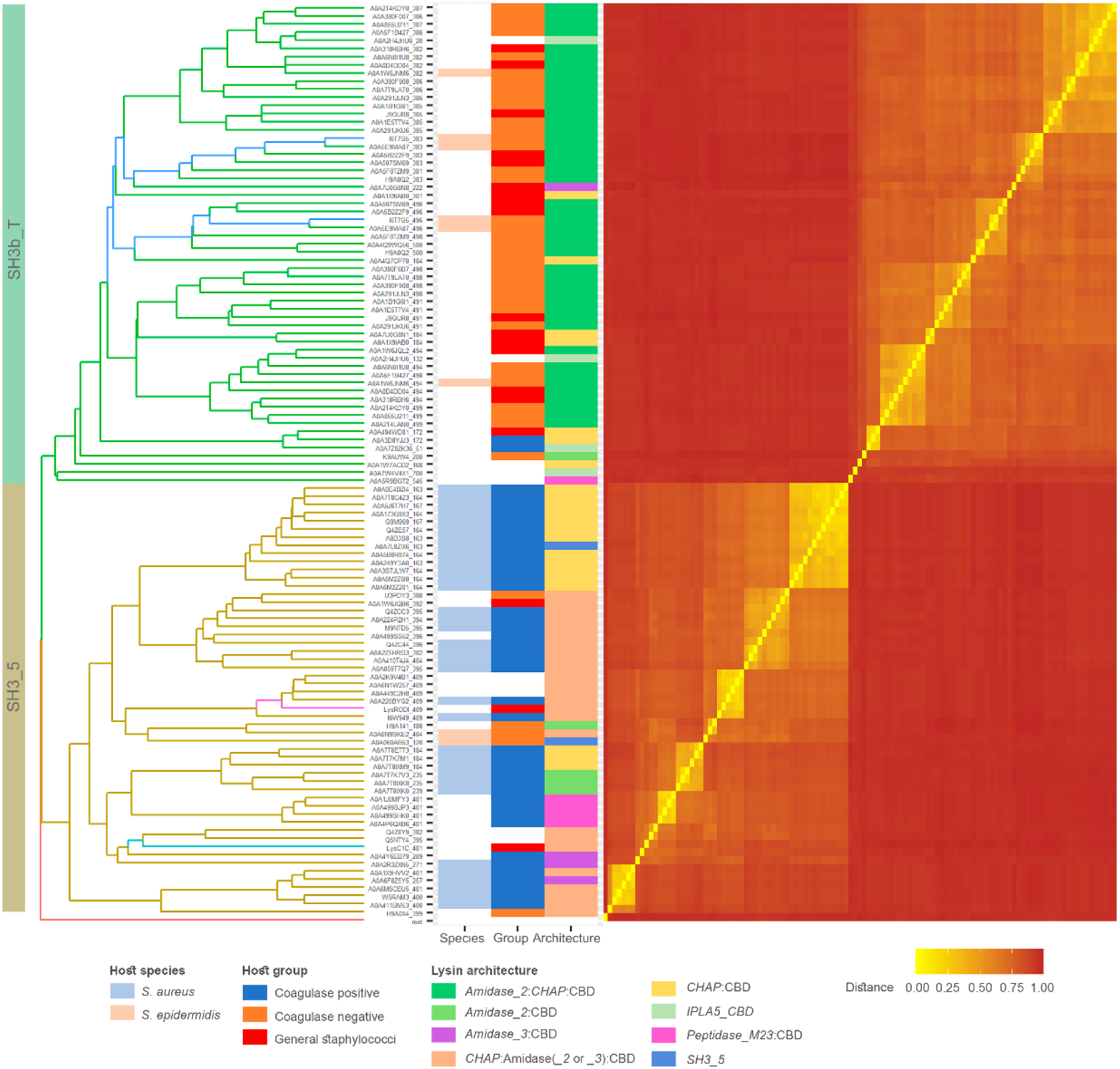
Sequence similarity analysis of a set of CBD sequences from either *SH3_5* or *SH3b_T* family. Branches of the UPGMA tree (left) based on the distance matrix (right) obtained from the MSA of the sequences are colored according to the CBD family, except those belonging to the endolysins of interest described in Figure 1, which are shown in a different color (IPLA5_CBD1 and IPLA5_CBD2 in dark blue, LysC1C in cyan, LysRODI in magenta). Tip labels are unique identifiers of each CBD comprising the UniProt accession number of the source protein plus the starting coordinate of the CBD delineation. Metadata associated to each CBD sequence is shown in the colored columns next to the tree.

### RODI_CBD, C1C_CBD and IPLA5_CBD present different binding profiles

To confirm the nature of IPLA5_CBD as a true CBD and explore the specificity differences to which the differential taxonomical distributions in **Figure 2** point, eGFP fusions of IPLA5_CBD, RODI_CBD and C1C_CBD were obtained. These eGFP-tagged domains were then assayed for binding capacity against a set of staphylococcal strains (**Figure 3a**). Well-marked trends could be observed in the binding specificity of the three different domains. RODI_CBD bound almost exclusively to *S. aureus* and a few other coagulase-negative staphylococci (CoNS), C1C_CBD was able to bind generally to all the tested staphylococci, and IPLA5_CBD bound preferentially to *S. epidermidis* plus other CoNS, although not in such a specific manner as RODI_CBD. A summary and the statistical significance supporting these trends are available in **Figure 3b**. Although generalizations should be made with care, these results provide an experimental explanation to the preferred association of *IPLA5_CBD* family to *S. epidermidis* and CoNS, while *SH3_5* is more commonly associated to *S. aureus* and only in some cases to CoNS or staphylococci in general (and LysC1C is an example of the latter). An additional conclusion of the results in **Figure 3** is that binding (*i.e.*, the magnitude of the recorded fluorescence value) seems to be generally lower when cells in stationary phase are used versus using exponential phase ones. An explanation for this may be provided by the nature of the ligand that has been described before for SH3b domains in anti-staphylococcal lysins, which has been usually identified as the peptidoglycan peptide moiety (22, 23, 43, 44). Stationary phase *S. aureus* cells are known to have fewer cross-links in the peptidoglycan (45), which would then mean a lower number of potential binding ligands for SH3b-like CBDs, thus explaining the results in **Figure 3**.

**Figure 3.**
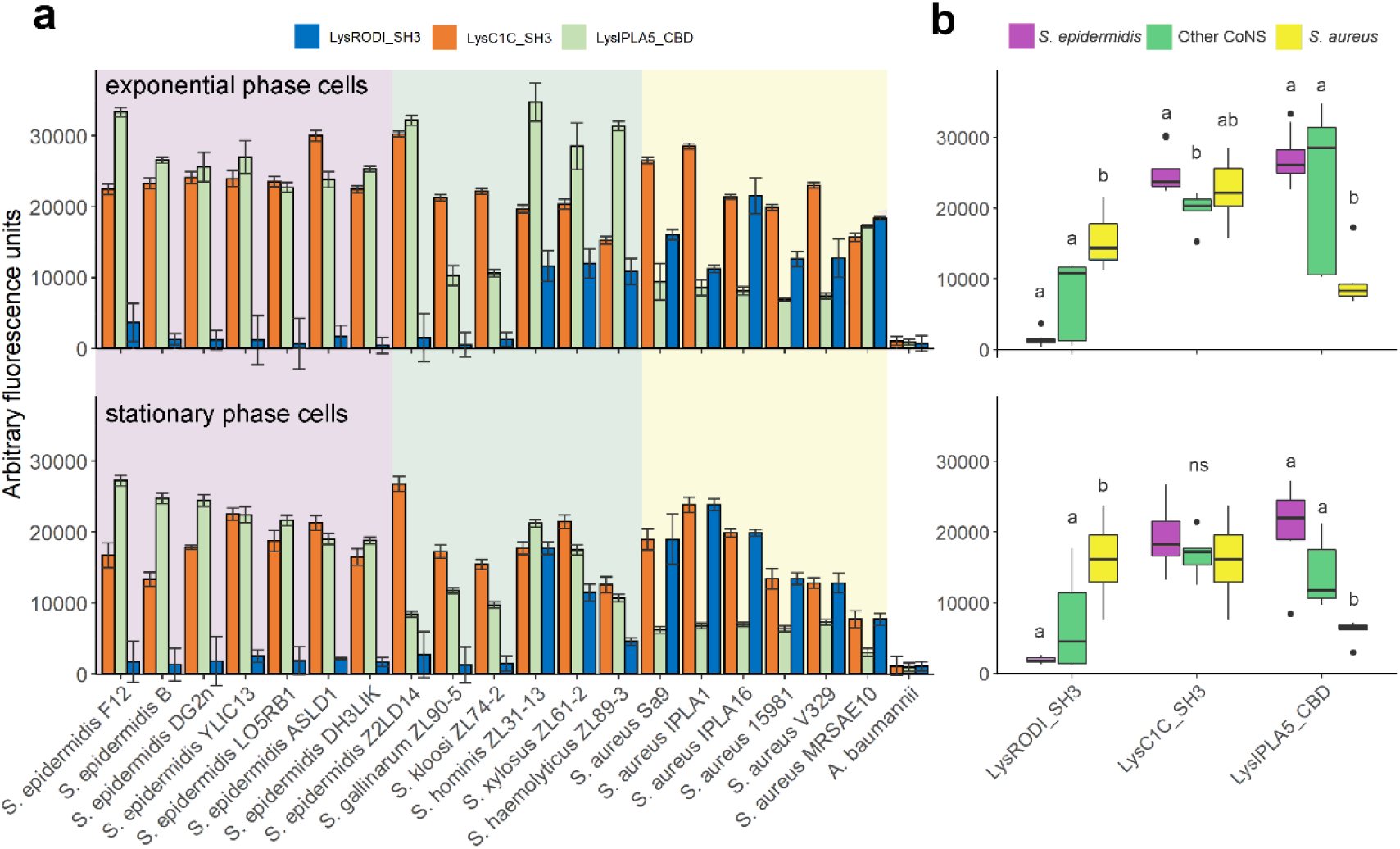
Binding specificity of CBDs from three anti-staphylococcal endolysins. (a) Binding capacity of RODI_CBD, C1C_CBD and IPLA5_CBD to a set of staphylococcal strains expressed as arbitrary fluorescence units. The background shades indicate the different bacterial species. (b) Summary and statistical comparison of the results in (a). Comparisons were made between results of each CBD class, grouped by staphylococcal strain (thus *S. epidermidis* vs. Other CoNS vs. *S. aureus* within each CBD). The Dunn test with Bonferroni adjustment was applied for multiple comparisons, and the significance of comparisons is depicted as small letters on top of each distribution box (different letters mean a statistically significant difference for a significance level of 0.05, ns means non-significant differences found within the class).

### The specificity profile of SH3b-like CBDs modulates the antibacterial spectrum of accompanying EADs

To better understand the contribution of the sole-CBD specificity profile to the full lysins’ activity spectrum, fusions of the *CHAP* EADs of LysRODI and LysC1C with either their wild-type CBD or IPLA5_CBD were obtained. Then, the MIC for each of the constructs, including only the EAD, was calculated against the full set of staphylococcal strains (**Figure 4**). A general conclusion from this experiment is that fusing a CBD normally improves the antimicrobial activity since all mean MIC log_2_ fold-change values in **Figure 4b** are negative, reflecting a decrease in the MIC as a result of fusing CBDs to the EADs. This is in accordance with the common notion that CBDs are necessary for the efficient action of endolysins. There are, however, a few cases in which the EAD-CBD fusion underperforms when compared with the sole EAD, namely RODI_CHAP is less active against some CoNS and a *S. epidermidis* strain when fused to RODI_CBD. In fact, the MIC improvement is not too impressive when fusing RODI_CBD to RODI_CHAP against any strain (with mean log_2_ fold-change values of about –1, in contrast with the 5 log_2_ fold decrease achieved by the IPLA5_CBD fusion against *S. epidermidis*, for example). This may be explained by the fact that RODI_CHAP already seems a highly optimized EAD against *S. aureus*, with clearly lower MIC values when compared with *S. epidermidis* or even the other coagulase negative staphylococci. The RODI_CHAP-IPLA5_CBD fusion does exhibit a remarkably lower MIC against *S. epidermidis* strains, but not against *S. aureus*, a behavior correlating to the specificity spectrum shown by IPLA5_CBD in **Figure 3**. In contrast with the *S. aureus*-specialized RODI_CHAP, C1C_CHAP is a broad-range EAD, as much as its native CBD is also broad range, although with a slight preference towards *S. epidermidis* (however non-significant). This intrinsic optimization towards *S. epidermidis* is more apparent when the effect of fusing C1C_CHAP to the different CBDs is considered: while, as expected, adding IPLA5_CBD improves the performance against *S. epidermidis*, the fusion with the broad-range C1C_CBD does not decrease the MIC value equally against *S. aureus* and *S. epidermidis*; in fact, it shows a similar effect to the IPLA5_CBD fusion (**Figure 4b**). This confirms a concomitant specialization of C1C_CHAP towards *S. epidermidis* peptidoglycan rather than that of *S. aureus*.

**Figure 4.**
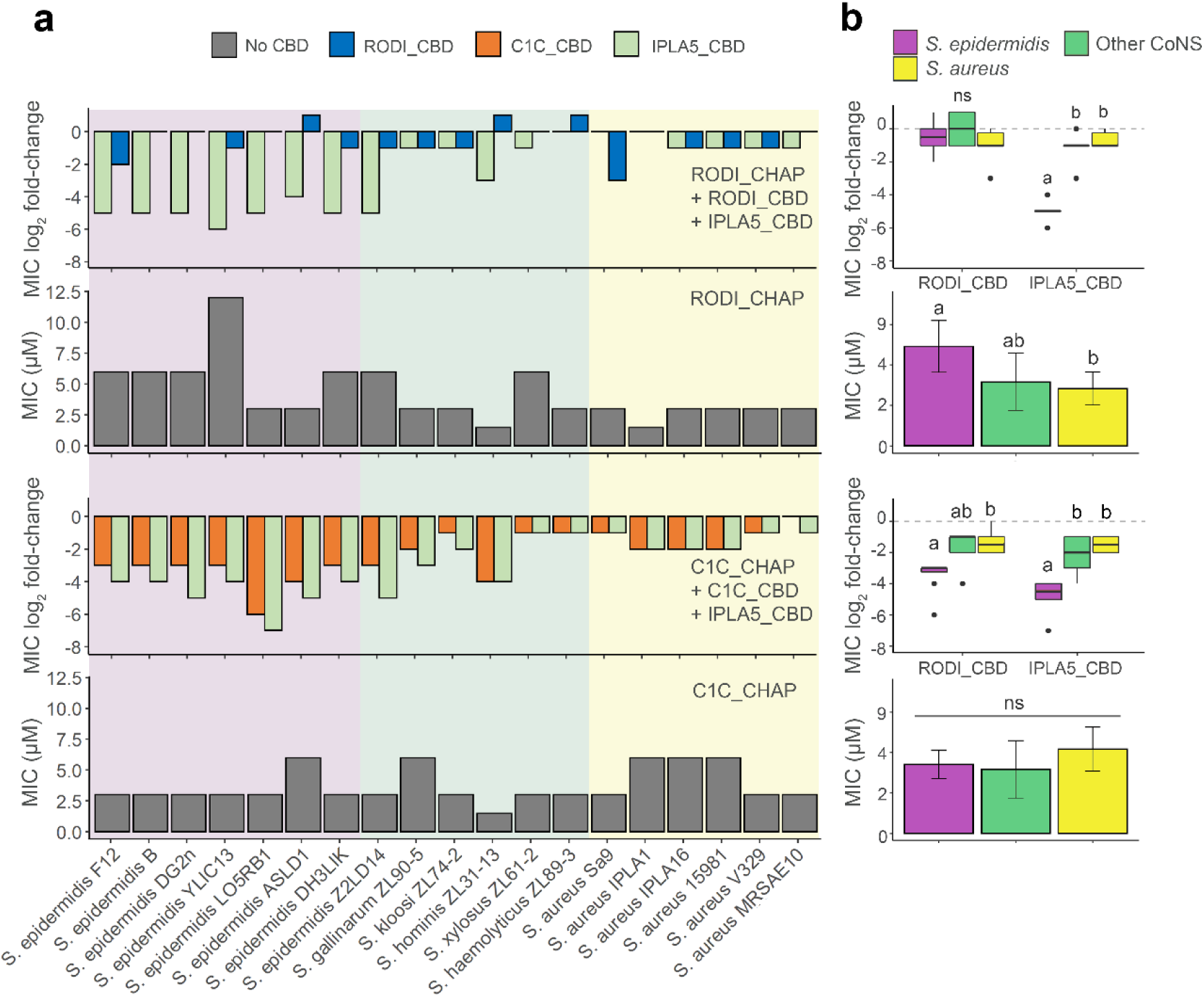
Impact of staphylococcal CBDs on the activity spectrum of EADs. (a) MIC values (grey bars) for the EADs RODI_CHAP (upper charts) and C1C_CHAP (lower charts) and log_2_ fold-change of the MIC for fusions of each of the EADs with their original CBD or with IPLA5_CBD with respect to the MIC of the EADs alone. Absolute MIC values can be consulted in **Supplementary Table S1**. The background shades indicate the different bacterial species. (b) Summaries of the data presented in (a) with the datapoints grouped by bacterial class (*S. epidermidis*, other CoNS or *S. aureus*). Statistically significant differences were tested between such bacterial groups using the Kruskal-Wallis test plus the Dunn test with Bonferroni adjustment for multiple comparisons. Boxplots or bars marked with different letters are significantly different to each other for a significance level of 0.05 (ns = non-significant difference).

### The difference in specificity between RODI_CBD and C1C_CBD could be explained by variability in key residues

The structures of the CBDs under investigation in this work were analyzed to find determinants for the perceived functional differences shown in **Figure 3** and **Figure 4**. To this end, MSAs were obtained using the representative sequence sets from **Figure 2**. A direct MSA-based comparison between the *SH3_5* and the *SH3b_T* domains was not possible due to their low reciprocal identity (**Figure 1c**); thus, the analysis was split to focus first on *SH3_5* domains (**Supplementary Figure S3**) and, therefore, on the structural differences that explain the different specificities of RODI_CBD and C1C_CBD. A set of key residues for peptidoglycan binding in *SH3_5* domains was determined from the thorough, previously published works on the *SH3_5* CBD of lysostaphin (22, 23). Then, the corresponding residues in RODI_CBD and C1C_CBD (**Table 3**) were identified assisted by MSA in **Supplementary Figure S3** and the comparison of the predicted 3D models of RODI_CBD and C1C_CBD with the experimentally determined structure of lysostaphin CBD (**Figure 5a**).

**Figure 5.**
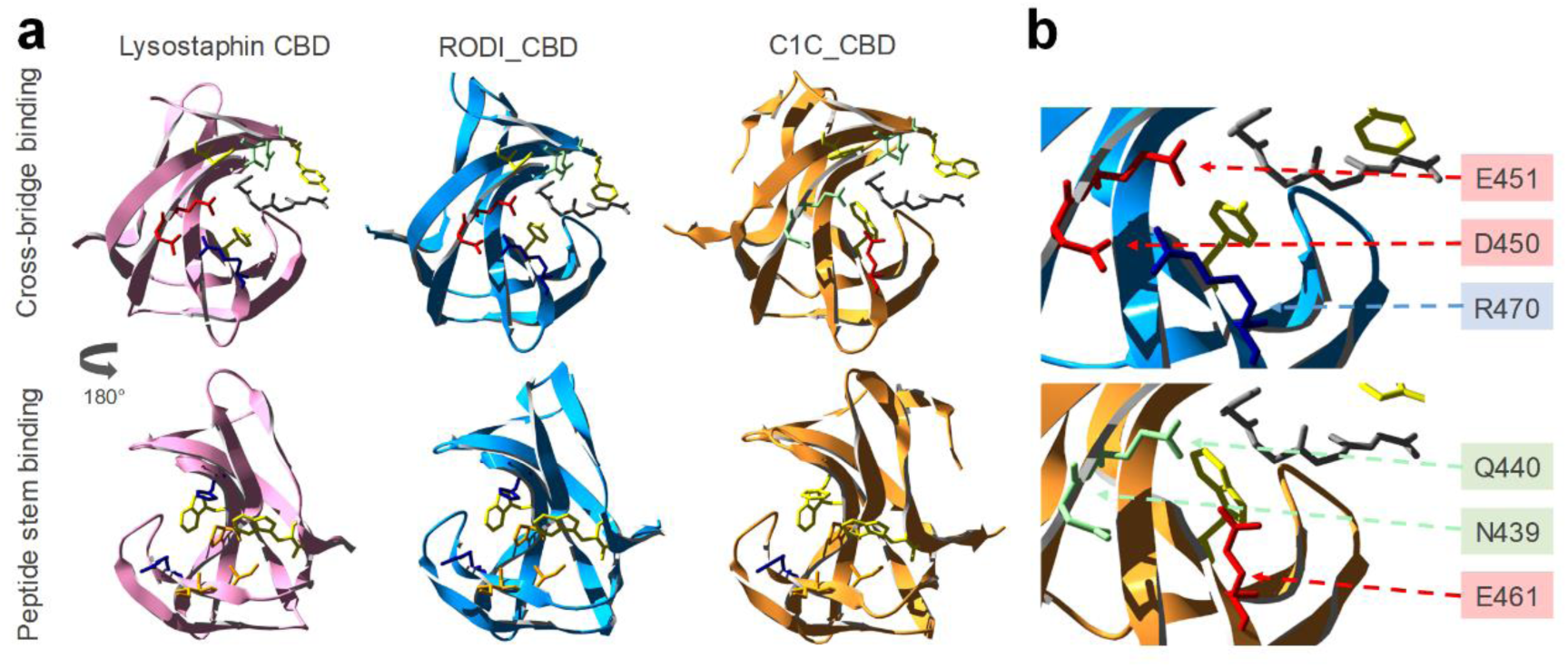
Structural comparison of lysostaphin CBD with those of LysRODI and LysC1C. (a) Crystal structure of lysostaphin (PDB 5LEO) compared with structural models of RODI_CBD (full protein pLDDT = 83.82) and C1C_CBD (full protein pLDDT = 86.67), displaying the key residues for binding the peptidoglycan cross-bridge and peptide stem as shown in **Table 3**. The grey chain represents a (Gly)_5_ peptide from PBD 5LEO, shown as a spatial reference for the cross-bridge binding pocket. (b) Detail of the cross-bridge-binding region of RODI_CBD and C1C_CBD and identification of the key differing residues between both domains. Amino acids are colored using the same code as described in **Table 3**.

**Table 3.** Key residues in *SH3_5* domains associated with binding the peptide stem or the cross-bridge according to (22, 23). Cells are colored to highlight different types of amino acids (green = polar non-charged, orange = aliphatic, yellow = aromatic, red = negatively charged, blue = positively charged). Residues are numbered according to the full sequence of the source proteins (lysostaphin = UniProt P10547).

| Lysostaphin | RODI_CBD | C1C_CBD |
| --- | --- | --- |
| **Cross-bridge binding** |  |  |
| N405 | N404 | N392 |
| Y407 | F406 | W394 |
| T409 | T408 | T396 |
| Y411 | Y410 | W398 |
| D450 | D450 | N439 |
| E451 | E451 | Q440 |
| R470 | R470 | E461 |
| Y472 | Y472 | W463 |
| **Peptide stem binding** |  |  |
| I424 | I424 | I413 |
| I425 | I425 | I414 |
| R427 | R427 | R416 |
| H458 | H458 | F449 |
| W460 | W460 | W451 |
| P474 | P474 | P465 |
| W489 | W491 | W479 |

**Table 3** shows little or no change in the chemical nature of the residues known to bind the peptide stem of the peptidoglycan across the three CBDs. Conversely, C1C_CBD displays radical shifts in amino acid properties at positions 439, 440 and 461, within the peptidoglycan cross-bridge binding site with respect to lysostaphin and RODI_CBD. Whereas C1C_CBD contains two polar, non-charged amino acids (N439, Q440) and a negatively charged one (E461), RODI_CBD and lysostaphin bear two negatively charged residues (D450, E451) and a positively charged one (R470) at the corresponding sites. Zooming into the analogous structures of these CBDs, it can be concluded that these changes imply a substantial rearrangement of the chemical environment at the groove that is assumed to bind the peptidoglycan cross-bridge. Particularly, D450 and R470 in RODI_CBD are at a distance between 2.98 Å and 4.13 Å (depending on the atoms considered), which makes it possible for them to form a salt bridge (46). This possibility is disrupted in C1C_CBD by the presence of N439 and E461 (**Figure 5b**).

The facts that (i) the key residues in RODI_CBD and lysostaphin CBD are relatively unchanged but (ii) there are obvious differences in C1C_CBD only at the cross-bridge-binding region is interpretable in the light of the experimental results presented in **Figure 3** and **Figure 4**. While RODI_CBD binds *S. aureus* specifically, as lysostaphin CBD does, C1C_CBD seems to have no clear preference between binding *S. aureus* or *S. epidermidis*. Assuming that the three *SH3_5* CBDs compared bind to the same ligand, the peptide moiety of peptidoglycan, which seems plausible given their sequence similarity (**Supplementary Figure S2**) and the conservation of the residues putatively devoted to binding the peptide stem (**Table 3**), then the differences found at the cross-bridge binding pocket of C1C_CBD must explain its promiscuous binding profile. In fact, the major difference between the peptidoglycans of *S. aureus* and *S. epidermidis* is the structure of their cross-bridges. While the cross-bridge of *S. aureus* is the well-known pentaglycine bridge, the cross-bridging peptide in *S. epidermidis* is either GGSGG or AGGGG (47). Since the introduction of a central serine residue in the cross-bridge is a known resistance mechanism to lysostaphin binding (48), it is reasonable that the RODI_CBD, structurally equivalent to lysostaphin CBD, is unable to bind the serine-containing *S. epidermidis* peptidoglycan cross-bridge. The specificity mechanism in lysostaphin CBD is thought to be one of steric constraint (the cross-bridge binding pocket can only accommodate a pentaglycine peptide (22)). Therefore, in C1C_CBD, the variants N394, Q440 and E461 should provide a greater flexibility for ligand placing at the cross-bridge binding site. This increased flexibility may be achieved by the disruption of the D450-R470 salt bridge in C1C_CBD (**Figure 5b)**.

### The IPLA5_CBD fold is closer to *PSA_CBD* or *PCD5_CBD* and may bind a different ligand than *SH3_5* CBDs

Given the low sequence similarity between the *SH3_5* and the *SH3b_T* sequences, a structure-based comparison was attempted using a few *bona fide* experimentally determined examples from the wider SH3b CBD superfamily (*i.e.*, lysostaphin CBD, PSA_CBD and PCD5_CBD), which were aligned to the predicted structure of a single IPLA5_CBD repeat (**Figure 6a**). This comparison initially suggested that IPLA5_CBD repeats were predicted in an SH3b fold more similar to that of the *PSA_CBD* family (PF18341) or the still poorly described *PCD5_CBD* family. When sets of representative sequences from *PSA_CBD* and *PCD5_CBD* families were added to a phylogenetic tree together with *SH3b_T* and *SH3_5*, this was made apparent by the clustering of the former three apart from the latter (**Figure 6b**).

**Figure 6.**
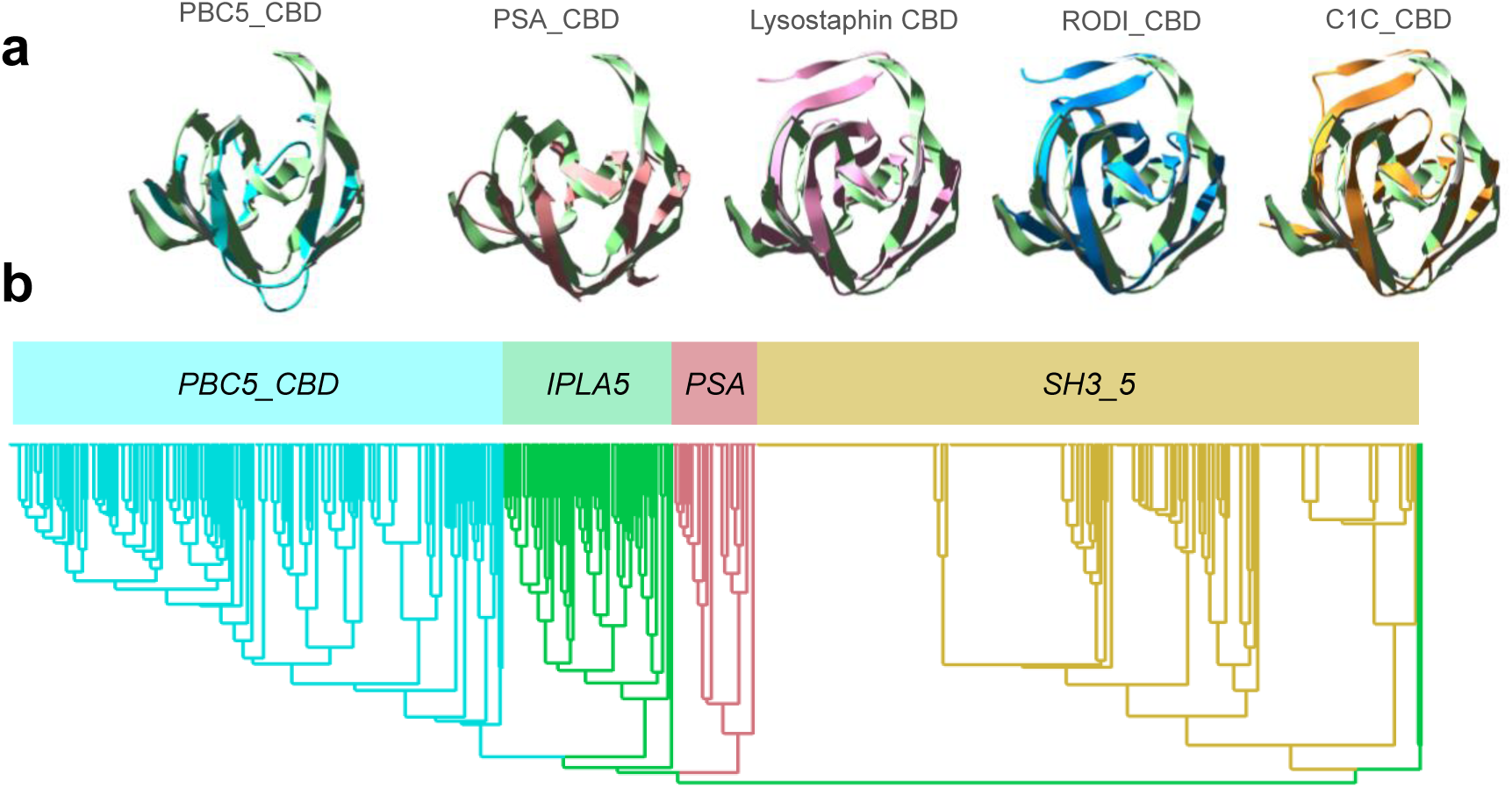
Contextualization of *SH3b_T* within the broader SH3b superfamily of CBDs. (a) Structural alignments of IPLA5_CBD first repeat (green) with lysostaphin CBD (PDB 5LEO, pink), the predicted structures of RODI_CBD (blue) and C1C_CBD (orange) and the first repeat from experimentally determined structures of CBDs from *PSA_CBD* and *PBC5_CBD* families (respectively, PDB 1XOV, pale red, and PDB 6ILU, cyan). (b) Phylogenetic tree comprising representatives from each of the aforementioned families (sequence data available in **Supplementary Dataset S1**).

A preliminary conclusion from this observation is that the ligand for *IPLA5_CBD* family may be different from that of *SH3_5* (*i.e.*, not the peptidoglycan cross-bridge or peptide moiety) given that *PSA_CBD* domains are known to bind sugar moieties in the teichoic acids of *Listeria* cells (25), while the ligand for PCD5_CBD has been described as the peptidoglycan glycan strands (26). Since *IPLA5_CBD* family is closer to *PSA_CBD* and *PCD5_CBD* than to *SH3_5*, it may bind a glycan moiety rather that a peptidic one. Given that the cell wall teichoic acids are also, in general, a differential feature between *S. aureus* and *S. epidermidis* (49), teichoic acids might be the ligand of IPLA5_CBD explaining its preference for *S. epidermidis* in detriment of *S. aureus*.

## DISCUSSION

In this work, we have identified a new family of SH3b-like domains that binds elements of the staphylococcal cell wall, with a preference towards *S. epidermidis*, and we have set a context for part of the underexplored diversity of the versatile SH3b-like folds among lysins. Here, we have compared four SH3b families present in phage endolysins: the *SH3_5* family commonly found in staphylococcal lysins, the newly described *SH3b_T* present in LysIPLA5, the listerial *PSA_CBD* and the *PBC5_CBD* domains found in *Bacillus* and their phages. All of them share the common β-barrel structure, typical of SH3 folds (**Figure 6a**), although with differing topologies (**Figure 7**). These differences may correlate with the type of cell wall ligands they recognize, namely (i) the peptidoglycan peptide moiety (or more specifically the cross-bridge) for *SH3_5* CBDs or (ii) different glycan moieties for *PSA_CBD*, *PBC5_CBD* and, perhaps, *IPLA5_CBD*. Regarding their topology, all of them conserve the general structure of the SH3 fold, and share the common SH3b trait of an extended RT β-hairpin (equivalent of the RT loop in SH3e, the topology of eukaryotic SH3 domains) plus the conserved central antiparallel β-sheets 2 to 4 (“β-core”, **Figure 7**). Their main topological differences are located (i) at the N- and C-proximal β-sheets, such as the presence/absence of an additional N-terminal β-sheet (β_0_, present in *SH3_5* and *SH3b_T*) or an additional β-sheet connecting β_4_ and β_5_ (β_4-5_, in *SH3_5* and *PSA_CBD*); and also (ii) at the RT β-hairpin, which is clearly more elongated in *PSA_CBD*, *PBC5_CBD* and *SH3b_T*, with the latter having the longest version. These differences should somehow account for the different ligands of the families, although this work does not provide concrete insights about the possible ligand.

**Figure 7.**
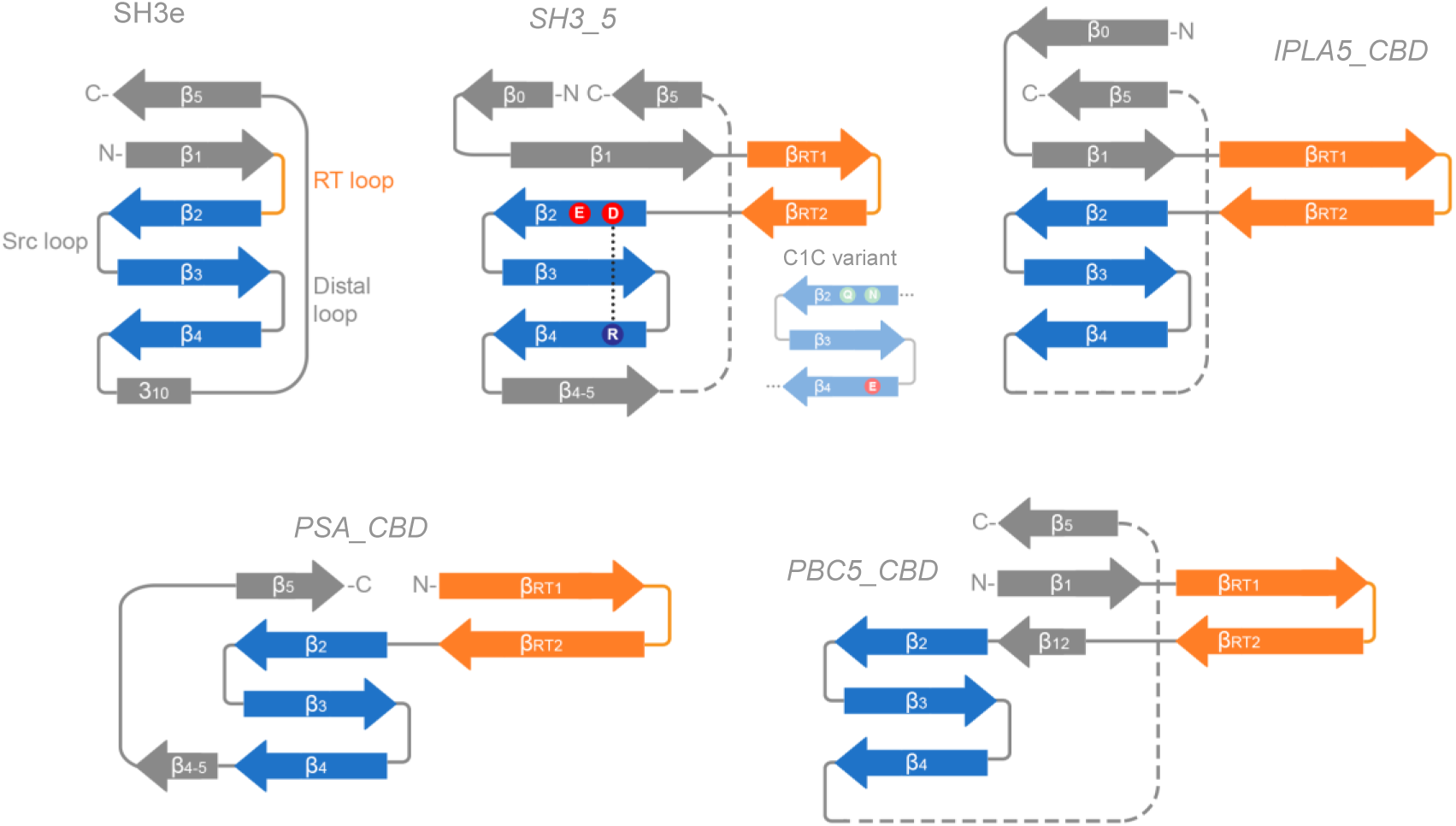
Compared topologies of SH3 domains. The RT loop in SH3e and its counterpart in SH3b families, the RT hairpin, is shown in orange, while the highly conserved β-core is in blue. Relevant residues at the β-core are displayed for the *SH3_5* family, including the D450-R470 salt bridge. β-sheets are numbered according to the canonical SH3 fold numeration, and extra sheets are given different names to facilitate the comparison.

On a different note, the results hereby presented show the versatility of the SH3b domains for evolving slightly different structures that bind different ligands, which is particularly evident in the comparison between RODI_CBD and C1C_CBD. While both belong to the same *SH3_5* family, their divergence only in a small set of key residues sets them rather radically apart in their specificity, determined by the nature of the peptidoglycan cross-bridge on the ligand side. This suggests that not only the SH3 fold can evolve towards topologically diverse families, each of them recognizing a different kind of ligand, but also within each family, SH3b domains can fine-tune the residues at the binding pockets to select the specific ligands they bind. Thus, the wide-spread presence of SH3b-like domains in endolysins from phages that infect very diverse groups of bacteria can be explained on the basis of this versatility of SH3 folds as “raw material” to evolve CBDs targeted at very specific ligands. However, the binding specificity dictated by CBDs, as we have shown, does not fully determine the activity spectrum of the derived lysins in which they are allocated (**Figure 4**). For example, the increased activity against *S. epidermidis* provided by IPLA5_CBD was more prominent when fused to an EAD with poor activity against this bacterium (RODI_CHAP) than when accompanying an EAD already optimized against *S. epidermidis* (C1C_CHAP). Thus, our set of results imply that, while the final antibacterial outcome correlates in terms of specificity to the observed specificity profile of the CBDs, it is also true that such outcome is influenced by the intrinsic activity range displayed by the EAD. Therefore, while the CBD can be a specificity determinant, it must be considered that it does not fully determine endolysin specificity, at least in the case of anti-staphylococcal lysins, and this may be of greater importance when engineering new tailor-made lysins.

## Acknowledgements

RV was supported by a postdoctoral fellowship of the ‘Bijzonder Onderzoeksfonds’ (BOF) Ghent University (01P10022). DGu was supported by the Research Foundation– Flanders (FWO) under grant G066919N. This work was also funded by MCIN/AEI/10.13039/501100011033/FEDER, UE, grant number PID2022-140988OB-I00 awarded to PG.

## Author contributions

YB, DGu, PG and AR conceived and designed research. DGu, DGr and RV conducted experiments. RV and DGu analyzed data. RV wrote the manuscript. All authors read and approved the manuscript.

## Statements and declarations

### Competing Interests

DGu is currently employed by Telum Therapeutics SL. DGr and YB are co-founders and (part-time) employee of Obulytix.

### Data availability

All data supporting the findings of this study are available within the paper and its Supplementary Information.

## References

1. Young R. 2014. Phage lysis: three steps, three choices, one outcome. J Microbiol 52:243–58.

2. Vázquez R, García E, García P. 2018. Phage lysins for fighting bacterial respiratory infections: a new generation of antimicrobials. Front Immunol 9:2252.

3. Dams D, Briers Y. 2019. Enzybiotics: enzyme-based antibacterials as therapeutics. Adv Exp Med Biol 1148:233–253.

4. Linden SB, Alreja AB, Nelson DC. 2021. Application of bacteriophage-derived endolysins to combat streptococcal disease: current state and perspectives. Curr Opin Biotechnol 68:213–220.

5. Murray CJL, Ikuta KS, Sharara F, Swetschinski L, Aguilar GR, Gray A, Han C, Bisignano C, Rao P, Wool E, Johnson SC, Browne AJ, Chipeta MG, Fell F, Hackett S, Haines-Woodhouse G, Hamadani BHK, Kumaran EAP, McManigal B, Achalapong S, Agarwal R, Akech S, Albertson S, Amuasi J, Andrews J, Aravkin A, Ashley E, Babin F-X, Bailey F, Baker S, Basnyat B, Bekker A, Bender R, Berkley JA, Bethou A, Bielicki J, Boonkasidecha S, Bukosia J, Carvalheiro C, Castañeda-Orjuela C, Chansamouth V, Chaurasia S, Chiurchiù S, Chowdhury F, Donatien RC, Cook AJ, Cooper B, Cressey TR, Criollo-Mora E, Cunningham M, Darboe S, Day NPJ, Luca MD, Dokova K, Dramowski A, Dunachie SJ, Bich TD, Eckmanns T, Eibach D, Emami A, Feasey N, Fisher-Pearson N, Forrest K, Garcia C, Garrett D, Gastmeier P, Giref AZ, Greer RC, Gupta V, Haller S, Haselbeck A, Hay SI, Holm M, Hopkins S, Hsia Y, Iregbu KC, Jacobs J, Jarovsky D, Javanmardi F, Jenney AWJ, Khorana M, Khusuwan S, Kissoon N, Kobeissi E, Kostyanev T, Krapp F, Krumkamp R, Kumar A, Kyu HH, Lim C, Lim K, Limmathurotsakul D, Loftus MJ, Lunn M, Ma J, Manoharan A, Marks F, May J, Mayxay M, Mturi N, Munera-Huertas T, Musicha P, Musila LA, Mussi-Pinhata MM, Naidu RN, Nakamura T, Nanavati R, Nangia S, Newton P, Ngoun C, Novotney A, Nwakanma D, Obiero CW, Ochoa TJ, Olivas-Martinez A, Olliaro P, Ooko E, Ortiz-Brizuela E, Ounchanum P, Pak GD, Paredes JL, Peleg AY, Perrone C, Phe T, Phommasone K, Plakkal N, Ponce-de-Leon A, Raad M, Ramdin T, Rattanavong S, Riddell A, Roberts T, Robotham JV, Roca A, Rosenthal VD, Rudd KE, Russell N, Sader HS, Saengchan W, Schnall J, Scott JAG, Seekaew S, Sharland M, Shivamallappa M, Sifuentes-Osornio J, Simpson AJ, Steenkeste N, Stewardson AJ, Stoeva T, Tasak N, Thaiprakong A, Thwaites G, Tigoi C, Turner C, Turner P, Doorn HR van, Velaphi S, Vongpradith A, Vongsouvath M, Vu H, Walsh T, Walson JL, Waner S, Wangrangsimakul T, Wannapinij P, Wozniak T, Sharma TEMWY, Yu KC, Zheng P, Sartorius B, Lopez AD, Stergachis A, Moore C, Dolecek C, Naghavi M. 2022. Global burden of bacterial antimicrobial resistance in 2019: a systematic analysis. The Lancet 399:629–655.

6. Jim O’Neill. 2016. Tackling drug-resistant infections globally: final report and recommendations. The Review on Antimicrobial Resistance, United Kingdom.

7. Tacconelli E, Carrara E, Savoldi A, Harbarth S, Mendelson M, Monnet DL, Pulcini C, Kahlmeter G, Kluytmans J, Carmeli Y, Ouellette M, Outterson K, Patel J, Cavaleri M, Cox EM, Houchens CR, Grayson ML, Hansen P, Singh N, Theuretzbacher U, Magrini N, WHO Pathogens Priority List Working Group. 2018. Discovery, research, and development of new antibiotics: the WHO priority list of antibiotic-resistant bacteria and tuberculosis. Lancet Infect Dis 18:318–327.

8. Abdelkader K, Gerstmans H, Saafan A, Dishisha T, Briers Y. 2019. The preclinical and clinical progress of bacteriophages and their lytic enzymes: the parts are easier than the whole. Viruses 11.

9. Fowler VG, Das AF, Lipka-Diamond J, Schuch R, Pomerantz R, Jauregui-Peredo L, Bressler A, Evans D, Moran GJ, Rupp ME, Wise R, Corey GR, Zervos M, Douglas PS, Cassino C. 2020. Exebacase for patients with *Staphylococcus aureus* bloodstream infection and endocarditis. J Clin Invest 130:3750–3760.

10. Fowler VG Jr, Das AF, Lipka-Diamond J, Ambler JE, Schuch R, Pomerantz R, Cassino C, Jáuregui-Peredo L, Moran GJ, Rupp ME, Lachiewicz AM, Kuti JL, Wise RA, Kaye KS, Zervos MJ, Nichols WG. 2024. Exebacase in Addition to Standard-of-Care Antibiotics for Staphylococcus aureus Bloodstream Infections and Right-Sided Infective Endocarditis: A Phase 3, Superiority-Design, Placebo-Controlled, Randomized Clinical Trial (DISRUPT). Clin Infect Dis ciae043.

11. Smug BJ, Szczepaniak K, Rocha EPC, Dunin-Horkawicz S, Mostowy RJ. 2023. Ongoing shuffling of protein fragments diversifies core viral functions linked to interactions with bacterial hosts. Nat Commun 14:7460.

12. Gerstmans H, Criel B, Briers Y. 2018. Synthetic biology of modular endolysins. Biotechnol Adv 36:624–640.

13. Vázquez R, García E, García P. 2021. Sequence-Function Relationships in Phage-Encoded Bacterial Cell Wall Lytic Enzymes and Their Implications for Phage-Derived Product Design. J Virol 95:e0032121.

14. Guillen D, Sanchez S, Rodriguez-Sanoja R. 2010. Carbohydrate-binding domains: multiplicity of biological roles. Appl Microbiol Biotechnol 85:1241–9.

15. Loessner MJ, Kramer K, Ebel F, Scherer S. 2002. C-terminal domains of Listeria monocytogenes bacteriophage murein hydrolases determine specific recognition and high-affinity binding to bacterial cell wall carbohydrates. Mol Microbiol 44:335–49.

16. Gutierrez D, Fernandez L, Rodriguez A, Garcia P. 2018. Are phage lytic proteins the secret weapon to kill *Staphylococcus aureus*? mBio 9.

17. Ikuta KS, Swetschinski LR, Aguilar GR, Sharara F, Mestrovic T, Gray AP, Weaver ND, Wool EE, Han C, Hayoon AG, Aali A, Abate SM, Abbasi-Kangevari M, Abbasi-Kangevari Z, Abd-Elsalam S, Abebe G, Abedi A, Abhari AP, Abidi H, Aboagye RG, Absalan A, Ali HA, Acuna JM, Adane TD, Addo IY, Adegboye OA, Adnan M, Adnani QES, Afzal MS, Afzal S, Aghdam ZB, Ahinkorah BO, Ahmad A, Ahmad AR, Ahmad R, Ahmad S, Ahmad S, Ahmadi S, Ahmed A, Ahmed H, Ahmed JQ, Rashid TA, Ajami M, Aji B, Akbarzadeh-Khiavi M, Akunna CJ, Hamad HA, Alahdab F, Al-Aly Z, Aldeyab MA, Aleman AV, Alhalaiqa FAN, Alhassan RK, Ali BA, Ali L, Ali SS, Alimohamadi Y, Alipour V, Alizadeh A, Aljunid SM, Allel K, Almustanyir S, Ameyaw EK, Amit AML, Anandavelane N, Ancuceanu R, Andrei CL, Andrei T, Anggraini D, Ansar A, Anyasodor AE, Arabloo J, Aravkin AY, Areda D, Aripov T, Artamonov AA, Arulappan J, Aruleba RT, Asaduzzaman M, Ashraf T, Athari SS, Atlaw D, Attia S, Ausloos M, Awoke T, Quintanilla BPA, Ayana TM, Azadnajafabad S, Jafari AA, B DB, Badar M, Badiye AD, Baghcheghi N, Bagherieh S, Baig AA, Banerjee I, Barac A, Bardhan M, Barone-Adesi F, Barqawi HJ, Barrow A, Baskaran P, Basu S, Batiha A-MM, Bedi N, Belete MA, Belgaumi UI, Bender RG, Bhandari B, Bhandari D, Bhardwaj P, Bhaskar S, Bhattacharyya K, Bhattarai S, Bitaraf S, Buonsenso D, Butt ZA, Santos FLC dos, Cai J, Calina D, Camargos P, Cámera LA, Cárdenas R, Cevik M, Chadwick J, Charan J, Chaurasia A, Ching PR, Choudhari SG, Chowdhury EK, Chowdhury FR, Chu D-T, Chukwu IS, Dadras O, Dagnaw FT, Dai X, Das S, Dastiridou A, Debela SA, Demisse FW, Demissie S, Dereje D, Derese M, Desai HD, Dessalegn FN, Dessalegni SAA, Desye B, Dhaduk K, Dhimal M, Dhingra S, Diao N, Diaz D, Djalalinia S, Dodangeh M, Dongarwar D, Dora BT, Dorostkar F, Dsouza HL, Dubljanin E, Dunachie SJ, Durojaiye OC, Edinur HA, Ejigu HB, Ekholuenetale M, Ekundayo TC, El-Abid H, Elhadi M, Elmonem MA, Emami A, Bain LE, Enyew DB, Erkhembayar R, Eshrati B, Etaee F, Fagbamigbe AF, Falahi S, Fallahzadeh A, Faraon EJA, Fatehizadeh A, Fekadu G, Fernandes JC, Ferrari A, Fetensa G, Filip I, Fischer F, Foroutan M, Gaal PA, Gadanya MA, Gaidhane AM, Ganesan B, Gebrehiwot M, Ghanbari R, Nour MG, Ghashghaee A, Gholamrezanezhad A, Gholizadeh A, Golechha M, Goleij P, Golinelli D, Goodridge A, Gunawardane DA, Guo Y, Gupta RD, Gupta S, Gupta VB, Gupta VK, Guta A, Habibzadeh P, Avval AH, Halwani R, Hanif A, Hannan MA, Harapan H, Hassan S, Hassankhani H, Hayat K, Heibati B, Heidari G, Heidari M, Heidari-Soureshjani R, Herteliu C, Heyi DZ, Hezam K, Hoogar P, Horita N, Hossain MM, Hosseinzadeh M, Hostiuc M, Hostiuc S, Hoveidamanesh S, Huang J, Hussain S, Hussein NR, Ibitoye SE, Ilesanmi OS, Ilic IM, Ilic MD, Imam MT, Immurana M, Inbaraj LR, Iradukunda A, Ismail NE, Iwu CCD, Iwu CJ, J LM, Jakovljevic M, Jamshidi E, Javaheri T, Javanmardi F, Javidnia J, Jayapal SK, Jayarajah U, Jebai R, Jha RP, Joo T, Joseph N, Joukar F, Jozwiak JJ, Kacimi SEO, Kadashetti V, Kalankesh LR, Kalhor R, Kamal VK, Kandel H, Kapoor N, Karkhah S, Kassa BG, Kassebaum NJ, Katoto PD, Keykhaei M, Khajuria H, Khan A, Khan IA, Khan M, Khan MN, Khan MA, Khatatbeh MM, Khater MM, Kashani HRK, Khubchandani J, Kim H, Kim MS, Kimokoti RW, Kissoon N, Kochhar S, Kompani F, Kosen S, Koul PA, Laxminarayana SLK, Lopez FK, Krishan K, Krishnamoorthy V, Kulkarni V, Kumar N, Kurmi OP, Kuttikkattu A, Kyu HH, Lal DK, Lám J, Landires I, Lasrado S, Lee S, Lenzi J, Lewycka S, Li S, Lim SS, Liu W, Lodha R, Loftus MJ, Lohiya A, Lorenzovici L, Lotfi M, Mahmoodpoor A, Mahmoud MA, Mahmoudi R, Majeed A, Majidpoor J, Makki A, Mamo GA, Manla Y, Martorell M, Matei CN, McManigal B, Nasab EM, Mehrotra R, Melese A, Mendoza-Cano O, Menezes RG, Mentis A-FA, Micha G, Michalek IM, Sá ACMGN de, Kostova NM, Mir SA, Mirghafourvand M, Mirmoeeni S, Mirrakhimov EM, Mirza-Aghazadeh-Attari M, Misganaw AS, Misganaw A, Misra S, Mohammadi E, Mohammadi M, Mohammadian-Hafshejani A, Mohammed S, Mohan S, Mohseni M, Mokdad AH, Momtazmanesh S, Monasta L, Moore CE, Moradi M, Sarabi MM, Morrison SD, Motaghinejad M, Isfahani HM, Khaneghah AM, Mousavi-Aghdas SA, Mubarik S, Mulita F, Mulu GBB, Munro SB, Muthupandian S, Nair TS, Naqvi AA, Narang H, Natto ZS, Naveed M, Nayak BP, Naz S, Negoi I, Nejadghaderi SA, Kandel SN, Ngwa CH, Niazi RK, Sá ATN de, Noroozi N, Nouraei H, Nowroozi A, Nuñez-Samudio V, Nutor JJ, Nzoputam CI, Nzoputam OJ, Oancea B, Obaidur RM, Ojha VA, Okekunle AP, Okonji OC, Olagunju AT, Olusanya BO, Bali AO, Omer E, Otstavnov N, Oumer B, A MP, Padubidri JR, Pakshir K, Palicz T, Pana A, Pardhan S, Paredes JL, Parekh U, Park E-C, Park S, Pathak A, Paudel R, Paudel U, Pawar S, Toroudi HP, Peng M, Pensato U, Pepito VCF, Pereira M, Peres MFP, Perico N, Petcu I-R, Piracha ZZ, Podder I, Pokhrel N, Poluru R, Postma MJ, Pourtaheri N, Prashant A, Qattea I, Rabiee M, Rabiee N, Radfar A, Raeghi S, Rafiei S, Raghav PR, Rahbarnia L, Rahimi-Movaghar V, Rahman M, Rahman MA, Rahmani AM, Rahmanian V, Ram P, Ranjha MMAN, Rao SJ, Rashidi M-M, Rasul A, Ratan ZA, Rawaf S, Rawassizadeh R, Razeghinia MS, Redwan EMM, Regasa MT, Remuzzi G, Reta MA, Rezaei N, Rezapour A, Riad A, Ripon RK, Rudd KE, Saddik B, Sadeghian S, Saeed U, Safaei M, Safary A, Safi SZ, Sahebazzamani M, Sahebkar A, Sahoo H, Salahi S, Salahi S, Salari H, Salehi S, Kafil HS, Samy AM, Sanadgol N, Sankararaman S, Sanmarchi F, Sathian B, Sawhney M, Saya GK, Senthilkumaran S, Seylani A, Shah PA, Shaikh MA, Shaker E, Shakhmardanov MZ, Sharew MM, Sharifi-Razavi A, Sharma P, Sheikhi RA, Sheikhy A, Shetty PH, Shigematsu M, Shin JI, Shirzad-Aski H, Shivakumar KM, Shobeiri P, Shorofi SA, Shrestha S, Sibhat MM, Sidemo NB, Sikder MK, Silva LMLR, Singh JA, Singh P, Singh S, Siraj MS, Siwal SS, Skryabin VY, Skryabina AA, Socea B, Solomon DD, Song Y, Sreeramareddy CT, Suleman M, Abdulkader RS, Sultana S, Szócska M, Tabatabaeizadeh S-A, Tabish M, Taheri M, Taki E, Tan K-K, Tandukar S, Tat NY, Tat VY, Tefera BN, Tefera YM, Temesgen G, Temsah M-H, Tharwat S, Thiyagarajan A, Tleyjeh II, Troeger CE, Umapathi KK, Upadhyay E, Tahbaz SV, Valdez PR, Eynde JV den, Doorn HR van, Vaziri S, Verras G-I, Viswanathan H, Vo B, Waris A, Wassie GT, Wickramasinghe ND, Yaghoubi S, Yahya GATY, Jabbari SHY, Yigit A, Yiğit V, Yon DK, Yonemoto N, Zahir M, Zaman BA, Zaman SB, Zangiabadian M, Zare I, Zastrozhin MS, Zhang Z-J, Zheng P, Zhong C, Zoladl M, Zumla A, Hay SI, Dolecek C, Sartorius B, Murray CJL, Naghavi M. 2022. Global mortality associated with 33 bacterial pathogens in 2019: a systematic analysis for the Global Burden of Disease Study 2019. The Lancet 400:2221–2248.

18. Becker K, Heilmann C, Peters G. 2014. Coagulase-negative staphylococci. Clin Microbiol Rev 27:870–926.

19. Oliveira H, Sampaio M, Melo LDR, Dias O, Pope WH, Hatfull GF, Azeredo J. 2019. Staphylococci phages display vast genomic diversity and evolutionary relationships. BMC Genomics 20:357.

20. Kamitori S, Yoshida H. 2015. Structure-Function Relationship of Bacterial SH3 Domains, p. 71–89. In Kurochkina, N (ed.), SH Domains: Structure, Mechanisms and Applications. Springer International Publishing, Cham.

21. Alvarez-Carreño C, Penev P, Petrov A, Williams L. 2021. Fold Evolution before LUCA: Common Ancestry of SH3 Domains and OB Domains. Mol Biol Evol 38.

22. Mitkowski P, Jagielska E, Nowak E, Bujnicki JM, Stefaniak F, Niedziałek D, Bochtler M, Sabała I. 2019. Structural bases of peptidoglycan recognition by lysostaphin SH3b domain. Sci Rep 9:5965.

23. Gonzalez-Delgado LS, Walters-Morgan H, Salamaga B, Robertson AJ, Hounslow AM, Jagielska E, Sabała I, Williamson MP, Lovering AL, Mesnage S. 2020. Two-site recognition of Staphylococcus aureus peptidoglycan by lysostaphin SH3b. Nat Chem Biol 16:24–30.

24. Beaussart A, Rolain T, Duchene MC, El-Kirat-Chatel S, Andre G, Hols P, Dufrene YF. 2013. Binding mechanism of the peptidoglycan hydrolase Acm2: low affinity, broad specificity. Biophys J 105:620–9.

25. Shen Y, Kalograiaki I, Prunotto A, Dunne M, Boulos S, I. Taylor NM, T. Sumrall E, R. Eugster M, Martin R, Julian-Rodero A, Gerber B, G. Leiman P, Menéndez M, Dal Peraro M, Javier Cañada F, J. Loessner M. 2021. Structural basis for recognition of bacterial cell wall teichoic acid by pseudo-symmetric SH3b-like repeats of a viral peptidoglycan hydrolase. Chem Sci 12:576–589.

26. Lee KO, Kong M, Kim I, Bai J, Cha S, Kim B, Ryu K-S, Ryu S, Suh J-Y. 2019. Structural Basis for Cell-Wall Recognition by Bacteriophage PBC5 Endolysin. Struct Lond Engl 1993 27:1355–1365.e4.

27. Gutiérrez D, Martínez B, Rodríguez A, García P. 2010. Isolation and Characterization of Bacteriophages Infecting Staphylococcus epidermidis. Curr Microbiol 61:601–608.

28. Gutiérrez D, Vandenheuvel D, Martínez B, Rodríguez A, Lavigne R, García P. 2015. Two Phages, phiIPLA-RODI and phiIPLA-C1C, Lyse Mono- and Dual-Species Staphylococcal Biofilms. Appl Environ Microbiol 81:3336–3348.

29. Dijkshoorn L, Aucken H, Gerner-Smidt P, Janssen P, Kaufmann ME, Garaizar J, Ursing J, Pitt TL. 1996. Comparison of outbreak and nonoutbreak Acinetobacter baumannii strains by genotypic and phenotypic methods. J Clin Microbiol 34:1519–1525.

30. Gerstmans H, Grimon D, Gutierrez D, Lood C, Rodriguez A, van Noort V, Lammertyn J, Lavigne R, Briers Y. 2020. A VersaTile-driven platform for rapid hit-to-lead development of engineered lysins. Sci Adv 6:eaaz1136.

31. Gutiérrez D, Garrido V, Fernández L, Portilla S, Rodríguez A, Grilló MJ, García P. 2020. Phage Lytic Protein LysRODI Prevents Staphylococcal Mastitis in Mice. Front Microbiol 11:7.

32. Potter SC, Luciani A, Eddy SR, Park Y, Lopez R, Finn RD. 2018. HMMER web server: 2018 update. Nucleic Acids Res 46:W200–W204.

33. Huang Y, Niu B, Gao Y, Fu L, Li W. 2010. CD-HIT Suite: a web server for clustering and comparing biological sequences. Bioinformatics 26:680–2.

34. Criel B, Taelman S, Van Criekinge W, Stock M, Briers Y. 2021. PhaLP: A Database for the Study of Phage Lytic Proteins and Their Evolution. Viruses 13.

35. Bodenhofer U, Bonatesta E, Horejš-Kainrath C, Hochreiter S. 2015. msa: an R package for multiple sequence alignment. Bioinformatics 31:3997–3999.

36. Zhou L, Feng T, Xu S, Gao F, Lam TT, Wang Q, Wu T, Huang H, Zhan L, Li L, Guan Y, Dai Z, Yu G. 2022. ggmsa: a visual exploration tool for multiple sequence alignment and associated data. Brief Bioinform 23:bbac222.

37. Charif D, Lobry JR. 2007. SeqinR 1.0-2: A Contributed Package to the R Project for Statistical Computing Devoted to Biological Sequences Retrieval and Analysis, p. 207–232. In Bastolla, U, Porto, M, Roman, HE, Vendruscolo, M (eds.), Structural Approaches to Sequence Evolution: Molecules, Networks, Populations. Springer, Berlin, Heidelberg.

38. Schliep KP. 2011. phangorn: phylogenetic analysis in R. Bioinformatics 27:592– 593.

39. Xu S, Li L, Luo X, Chen M, Tang W, Zhan L, Dai Z, Lam TT, Guan Y, Yu G. 2022. Ggtree: A serialized data object for visualization of a phylogenetic tree and annotation data. iMeta 1:e56.

40. Mirdita M, Schütze K, Moriwaki Y, Heo L, Ovchinnikov S, Steinegger M. 2022. ColabFold: making protein folding accessible to all. 6. Nat Methods 19:679–682.

41. Guex N, Peitsch MC, Schwede T. 2009. Automated comparative protein structure modeling with SWISS-MODEL and Swiss-PdbViewer: a historical perspective. Electrophoresis 30 Suppl 1:S162–173.

42. Wickham H. 2016. ggplot2 elegant graphics for data analysis, 2nd ed. Springer International Publishing.

43. Grundling A, Schneewind O. 2006. Cross-linked peptidoglycan mediates lysostaphin binding to the cell wall envelope of *Staphylococcus aureus*. J Bacteriol 188:2463–72.

44. Lu JZ, Fujiwara T, Komatsuzawa H, Sugai M, Sakon J. 2006. Cell wall-targeting domain of glycylglycine endopeptidase distinguishes among peptidoglycan cross-bridges. J Biol Chem 281:549–58.

45. Zhou X, Cegelski L. 2012. Nutrient-dependent structural changes in S. aureus peptidoglycan revealed by solid-state NMR spectroscopy. Biochemistry 51:8143– 8153.

46. Kumar S, Nussinov R. 2002. Close-Range Electrostatic Interactions in Proteins. ChemBioChem 3:604–617.

47. Schleifer KH, Kandler O. 1972. Peptidoglycan types of bacterial cell walls and their taxonomic implications. Bacteriol Rev 36:407–77.

48. Batool N, Ko KS, Chaurasia AK, Kim KK. 2020. Functional Identification of Serine Hydroxymethyltransferase as a Key Gene Involved in Lysostaphin Resistance and Virulence Potential of Staphylococcus aureus Strains. 23. Int J Mol Sci 21:9135.

49. Du X, Larsen J, Li M, Walter A, Slavetinsky C, Both A, Sanchez Carballo PM, Stegger M, Lehmann E, Liu Y, Liu J, Slavetinsky J, Duda KA, Krismer B, Heilbronner S, Weidenmaier C, Mayer C, Rohde H, Winstel V, Peschel A. 2021. Staphylococcus epidermidis clones express Staphylococcus aureus-type wall teichoic acid to shift from a commensal to pathogen lifestyle. Nat Microbiol 6:757–768.

